# Desmoplakin mutations in cardiac fibroblasts cause TGFβ1-mediated pathological fibrogenesis in desmoplakin cardiomyopathy via beclin-1 regulation

**DOI:** 10.1101/2024.09.09.612149

**Authors:** Chuanyu Wei, Shing-Fai Chan, Ardan M. Saguner, Corinna Brunckhorst, Firat Duru, Joseph E. Marine, Cynthia A. James, Hugh Calkins, Daniel P. Judge, Weinian Shou, Huei-Sheng Vincent Chen

## Abstract

**Background:** Pathological fibrosis is a major finding in cardiovascular diseases and can result in arrhythmia and heart failure. Desmosome gene mutations can lead to arrhythmogenic cardiomyopathy (ACM). Among ACM, pathogenic desmoplakin (*DSP*) variants cause a distinctive cardiomyopathy with excessive cardiac fibrosis that could precede ventricular dysfunction. *DSP* variants are also linked to other fibrotic diseases. Whether DSP plays any role in pathological fibrosis remain unknown.

**Methods:** Mesenchymal stromal cells (MSCs) are resident fibroblast-like cells that are responsible for fibrogenesis in most organs, including hearts. We first used unbiased genome-wide analyses to generate cardiac fibroblasts-like, induced pluripotent stem cell-derived MSCs from normal donors and ACM patients with *DSP* mutations. We then studied the fibrogenic responses of cardiac MSCs to transforming growth factor beta-1 (TGF-β1) using Western/Co-IP, autophagy assay, gene knockdowns/over-expressions, genomic analyses, mouse DSP knockdown models, immunostaining, and qPCR.

**Results:** TGFβ1 induced excessive accumulations of vimentin (VIM)/fibrillar collagens, and over-activated fibrotic genes in *DSP-*mutant MSCs when compared to normal MSCs. In normal MSCs, VIMs bind to wild-type DSP during normal fibrogenesis after TGFβ1. *DSP-*mutant MSCs exhibited a haplo-insufficient phenotype with increased DSP-unbound VIMs that sequestered beclin-1 (BECN1) from activating autophagy and caveolin-1 (CAV1)-mediated endocytosis. Decreased autophagy caused collagen accumulations and diminished CAV1 endocytosis resulted in abnormal CAV1 plaque formation that over-activated fibrotic genes [*COL1A1, COL3A1,* and fibronectin (*FN*)] via heightened p38 activities after TGFβ1. Genome-wide analysis and DSP knockdown in mouse fibroblasts confirmed this novel role of *DSP* mutations in pathological fibrosis. Overexpression of VIM-binding domains of DSP could suppress pathological fibrosis by increasing collagen autophagic degradation and decreasing fibrotic gene expressions.

**Conclusions:** Our data reveal that DSP deficiency in MSCs/fibroblasts leads to exaggerated fibrogenesis in DSP-cardiomyopathy by decreasing BECN1 availability for autophagy and CAV1-endocytosis. Overexpression of VIM binding domains of DSP could be a new strategy to treat pathological fibrosis.

## Introduction

Arrhythmogenic Cardiomyopathy (ACM) is an inherited cardiomyopathy frequently caused by pathogenic variants in five desmosomal genes, including plakophilin-2 (*PKP2*), desmoplakin (*DSP*), desmoglein 2, desmocollin 2 (*DSC2*), and plakoglobin. Progressive fibro-fatty replacement of cardiomyocytes (CMs) with increased CM apoptosis is the pathological hallmark of ACM, which may cause sudden death in the young.^1–4^ Among desmosomal mutations, human ACM hearts with *DSP* mutations are different from other types of desmosomal mutations by displaying more pathological fibrosis than fatty infiltration (adipogenesis) and affecting both ventricles.^5,6^ *DSP* knockout or decreased *DSP* expression has also been implicated in the pathogenesis of fibrotic diseases, such as idiopathic pulmonary fibrosis (IPF)^7^ and hypertrophic scars (a form of pathological fibrosis).^8^ Whether and how *DSP* mutations could cause or mediate pathological fibrosis remain to be determined.

Pathological cardiac fibrosis is usually characterized by excessive production and accumulation of cytosolic vimentin (VIM) and extracellular matrices (ECM, particularly fibrillar collagens) in diseased hearts^9–12^ and may lead to lethal ventricular arrhythmia and heart failure. Resident cardiac fibroblasts are generally considered the principal source of cardiac fibrosis.^13^ However, cardiac fibroblasts display heterogeneity^14^ and lack commonly accepted markers for identification.^15^ In contrast, mesenchymal stromal/stem cells (MSCs), morphologically indistinguishable from fibroblasts, have internationally defined characteristics,^16^ can be isolated reproducibly from various organs, and are known adult resident progenitor cells for myofibroblasts (MBs, activated fibroblasts)^13^ that produce fibrosis in most organs,^17^ including the heart. However, obtaining cardiac MSCs or fibroblasts from cardiac patients is rarely performed. Recent advances in reprogramming somatic cells into induced pluripotent stem cells (iPSCs) have enabled the derivation of MSCs from iPSCs (iPSC-MSCs);^18,19^ this process exhibits high purity/yield of MSCs and does not require any invasive procedure. We primarily used an iPSC-MSC derivation protocol^19^ that drove pluripotent iPSC through an early mesoderm stage before the final derivation of MSCs, simulating cardiac development *in vivo*.

We used transforming growth factor beta-1 (TGFβ1), a well-known fibrosis activator,^20^ to induce fibrogenic responses in cardiac MSCs because TGFβ1 level was particularly elevated in ACM mouse hearts^21^ and in human ACM patients.^22^ In general, fibrogenic responses to TGFβ1 in MSCs could be reparative or maladaptive. In normal reparative responses, TGFβ1 induces transient activation of fibrogenic genes and proteins (e. g. collagens) after injury for physiological repair,^23^ which processes are controlled and later terminated by TGFβ1-induced endocytosis-lysosome (that terminate TGFβ1 signaling)^23,24^ and autophagy (that degrades redundant collagens)^25^ pathways. In contrast, the balance between activation and termination of the TGFβ pathway appears lost in maladaptive responses to TGFβ1, resulting in prolonged and excessive production of ECM that contribute to pathological organ fibrosis.^10,22–26^

Here, we used unbiased RNA sequencing (RNAseq) to show that our iPSC-MSCs are similar to cardiac fibroblasts grown directly from the source human heart by genome-wide profiling. This is the first time that cardiac fibroblast-like iPSC-MSCs are characterized by genome-wide comparisons with fibroblasts directly grown from the same source donor heart rather than using a few highly selective markers. We then generated cardiac iPSC-MSCs from healthy donors and 2 unrelated ACM patients with pathogenic heterozygous *DSP* variants to study how *DSP* mutations affect their fibrogenic potentials. We found that *DSP-*mutant MSCs had no mutant but ∼50% wild-type (WT) DSP proteins (a DSP-deficient and haploinsufficiency phenotype). Importantly, TGFβ1 induced excessive accumulations of VIMs/fibrillar collagens and overactivated fibrotic genes [*COL1A1, COL3A1,* and fibronectin (*FN*)] in *DSP-*mutant MSCs. In normal MSCs, VIMs bound to WT DSPs during normal fibrogenesis, yet in *DSP*-deficient MSCs, decreased WT DSPs led to increased DSP-unbound VIMs that sequestered BECN1 and decreased its availability for autophagy/lysosome degradation of collagens. Decreased BECN1 availability also reduced caveolin-1 (CAV1)-mediated endocytosis, leading to abnormal formation of long CAV1 plaques that overactivated fibrotic genes via heightened p38 activities. *DSP* knockdown in normal MSCs and over-expression of *DSP* in mutant MSCs further support that *DSP* mutations lead to DSP deficiency, which causes pathological fibrogenesis via deregulated BECN1-VIM interactions. We also generated mice with heterozygous deletion of *DSP* in activated fibroblasts, which showed aggressive cardiac fibrosis to TGFb1 *in vivo* via the same pathogenic mechanisms. Furthermore, genome-wide gene expression analysis validated this novel pathogenic role of DSP deficiency in pathological fibrosis. Most importantly, over-expression of VIM-binding domains of DSP suppressed the pathological fibrosis in *DSP*-mutant MSCs, which could be a new strategy to treat pathological cardiac fibrosis.

## Methods

### Data availability

All data associated with this study and all unprocessed blots are included in the Supplemental Materials.

### Heart tissues and generation of human iPSCs

The generation and characterization methods of normal iPSC (SW) and desmosomal mutant iPSC lines were published previously.^27^ We obtained *DSP* mutant human peripheral blood mononuclear cells (PBMCs) from Johns-Hopkins University (JHU) and University of California-San Diego (UCSD) with approved protocols by relevant institutional review boards (IRBs) at each institution. All living human participants gave written informed consent. Frozen human normal ventricular tissues from a 38 years old donor heart without known heart disease were provided by AnaBios Corp. Frozen ventricular tissues from an ACM patient with a *DSP* (c.4531G>T, p.Q1511X) mutation were provided by University of Zurich under approved study protocols. PBMCs were reprogrammed using episomal vectors containing Oct4, Sox2, c-Myc and Klf4 to generate integration-free iPSC colonies according to published episomal methods (see Supplemental Methods for details). Human iPSCs and H9 human embryonic stem cells (hESCs) were maintained on a feeder-free system using Matrigel coated culture plates and TeSR-E8 media Kits (STEMCELL Technology Inc.).^27^ All culture media and reagents were purchased from Invitrogen (Carlsbad, CA)/Thermo Fisher Scientific (Waltham, MA) unless indicated otherwise.

### Generation of human normal and *DSP*-mutant iPSC-MSCs

We used a well-establish protocol to generate iPSC-MSCs via an early mesoderm intermediate stage as previously described^19^ to simulate cardiac development *in vivo* (see Supplemental Methods for details). We plated harvested iPSC-MSCs on non-coasted plastic 12-well plates and only fibroblasts-like cells attached to plastic surfaces in 20 minutes after plating were selected (plastic adhesion selection), which enabled the isolation and purification of iPSC-derived fibroblasts that fulfilled published criteria for MSCs.^16^

### Culturing fibroblasts from human and mouse tissues

A small piece (1-2 mm^2^) of unused and redundant skin or heart tissues from a human subject (SW), or from ventricular tissues of various mouse hearts was minced and placed into one well of 6-well non-coated plate and cultured in 3 ml MSC medium with Normocin (Invitrogen) and Gentamicin (GIBCO).^28^ The media was changed daily for 3 days and then every 2-3 days. Tissue pieces and nonadherent cells were removed in 7-10 days (in 3-4 days for mouse tissues) when fibroblasts appeared in the well. The fibroblasts were cultured to 90-95% confluence (Passage 0) and were then harvested by trypsin for further expansion or storage. Cardiac fibroblasts at passages ³ 3 were used for the experiments when sufficient amounts of fibroblasts were obtained. Human skin fibroblasts required more passages to yield sufficient amounts of fibroblasts for research.

### MSC characterization using flow cytometry

The MSCs were prepared and characterized according to the standard flow cytometry protocol at Indiana University Flow Cytometry Core (see Supplemental Methods for details). Antibodies used are listed in Table S1.

### Karyotype analysis

Karyotype analysis was performed by Cell Line Genetics (Madison, WI).

### Total mRNA isolation and quantitative real-time PCR (qPCR)

Total RNA from cultured cells was prepared using mirVana miRNA Isolation Kit (Invitrogen) according to the manufacturer’s instruction. For mRNA qPCR, about 200 ng of RNA was then reverse transcribed to complementary DNA (cDNA) using Quantitect Reverse Transcription Kit (Qiagen) according to the manufacturer’s protocol. qPCR was performed with 400 ng cDNA for 40 cycles using SYBR Green I (Roche) and Roche LightCycler 96 according to protocols recommended by the manufacturer. Primer sequences used are listed in Table S2. Quantification for mRNA expression levels was carried out by correcting for amplification efficiency of the primer using a standard curve, followed by normalizing transcript levels to the amount of total ubiquitously-expressed GAPDH transcripts. Relative expression levels of mRNA were calculated using the 2^-ΔΔCt^ method. 2-3 physical replicates of each sample from ≥ 3 independent sets of experiments (biological replicates) were performed. To construct each histogram, mRNA levels were normalized to the baseline mRNA levels of normal MSCs (as 1 or 100%) in all figures unless indicated otherwise.

### Genomic and cDNA sequence analysis

Genomic DNA at 200 ng per PCR reaction from undifferentiated iPSCs was collected using QIAamp DNA Mini Kit (Qiagen Cat. # 51304) and PCR reaction was set up using DreamTag Green PCR Master Mix (Thermo Scientific Cat. # K1081) following the manufacturer’s instructions. PCR products were prepared and sent to either GenScript (Piscataway, NJ) or Genewiz (South Plainfield, NJ) for DNA sequencing services according to each company’s standard requirement. Sequences of primers used are listed in Table S3.

### Immunoblotting (western blotting), immunoprecipitation (IP), and Co-IP

Standard techniques were used for western blots, IP, and Co-IP (see Supplemental Methods for details). Antibodies used with indicated dilutions in 5 % BSA are listed in Table S4. DSP antibody [Abcam #ab71690, stains both DSP-I (∼332 kD) and DSP-II (∼260 kD) isoforms]^29,30^ was used for all western blots in this study except the sub-Figure S2E.

### Sircol Collagen Assay

To examine collagen released into the cell culture media, normal and mutant MSCs were grown in 12-well plates with or without 1 ng/mL TGF-β1 for 72 hrs in the absence or presence of specific drugs. The supernatant was then collected and collagen content was measured by using the Sircol Assay according to the manufacturer’s instructions (Biocolor; Cat. No. S1000).

### Immunocytochemistry (IHC) on cells and human heart tissues

Standard techniques were used for IHC on cultured cells and cryo-preserved human heart tissues (see Supplemental Methods for details). Primary and secondary antibodies used are listed in Table S5. Alexa Fluor 488 and Alexa Fluor 594 anti-mouse or anti-rabbit IgG (Invitrogen) were the secondary antibodies used for the fluorescence imaging. Samples were imaged using confocal microscopy (Leica confocal). CAV1 plaque lengths, cell sizes, and CAV1-TGFβ3 receptor overlap lengths were measured using Image J software.

### Imaging autophagy with Premo™ autophagy tandem sensor RFP-GFP-LC3B

The MSCs (2 × 10^5^ cells/well) were cultured in a Nunc™ Lab-Tek 8 Chamber Slide System and allowed to adhere overnight. Various stages of autophagy were monitored using Premo™ Autophagy Tandem Sensor RFP-GFP LC3B Kit (Life Technologies) according to the manufacturer’s instructions (see Supplemental Methods for details).

### Plasmids and siRNAs

We used PCR and DSP isoform-I-GFP vector (Addgene plasmid#32227) to generate the HA-DSPΔN fragment (corresponding to amino acids 1057-2871) and subcloned this fragment to pcDNA3.1 vector so as to over-express HA-DSPΔN in mutant MSCs (see Supplemental Methods for details). Wildtype full-length beclin-1 cDNA (pcDNA4-beclin-1, Addgene plasmids#24388) was used for plasmid transfection and was cloned into pCDH-CMV-MCS-EF1-GFP lentivector (System Biosciences, Inc.). The ON-TARGET siRNA-SMART pool L-003551-00-0005 or L-003467-00-0005 was used for *VIM* and *CAV1* knockdown (Horizon Discovery), respectively. A pool of four siRNA duplexes against *DSP* was used (Horizon Discovery) to knock down *DSP*. We used AllStars Negative Control siRNA (Qiagen) as the control nonsilencing siRNA (siCTR).

### Transfection of plasmids

Plasmid cDNAs of wildtype *BECN1*, *BECN1*-GFP, *HA-DSPΔN*, or empty vector were transfected by using Lipofectamine 2000 in accordance with the manufacture’s instruction.

### RNA interference/transfection of synthetic siRNAs

MSCs were cultured in antibiotics-free MSC medium to 70–80% confluency, and then synthetic siRNA targeting human *VIM, CAV1* or *DSP*, and AllStars Negative Control siRNA at 50 nM were transfected into MSCs using Lipofectamine RNAiMAX Transfection Reagent (Thermo Fisher Scientific) following the manufacturer’s instructions. At 48 hours after the first transfection, a second transfection of siRNA targeting each gene at 50 nM was performed to obtain the best knockdown efficiency. After incubation for additional 48 hours, total protein was extracted to confirm the decreased proteins levels of each target gene by western blots.

### Electroporation of MSCs

To evaluate the effects of *BECN1* or *HA-DSPΔN* OE on mRNA and protein levels, electroporation was performed using the Neon Transfection System 100 µL Kit (Thermo Fisher Scientific) according to the manufacturer’s instructions. Electroporation was performed using parameters to obtain the best cell survival and transfection efficiency: voltage 990-1200 V; interval 20-40 ms; 1-2 pulses. After electroporation, the cell suspension was transferred and cultured in antibiotics-free MSC media.

### Generation and study of mice with heterozygous *DSP* deletion in myofibroblasts

In house mice with tamoxifen-inducible, periostin-specific cre recombinase (*POSTN^MRM^*-Cre+) genotypes^31^ were intercrossed with mice with heterozygous *DSP*-floxed mice (*DSP^w/f^*, Jax #026665, The Jackson Lab). Mice with *POSTN^MCM^*-Cre+; *DSP^w/f^* and P*OSTN^MCM^*-Cre-; *DSP^w/f^* genotypes were selected. We injected tamoxifen intraperitoneally (i. p.) for 5 days starting at postnatal 4 weeks of age to all mice but tamoxifen would only induce myofibroblast-specific heterozygous deletion of *DSP* in *POSTN^MCM^*-Cre+ mice. We then injected 1 µg TGFβ1 daily by i. p. for 7-8 days to all 6-week-old mice and mouse hearts were harvested for research after completing TGFβ1 injection. Sirius Red-fast green staining for collagens was performed as described previously^21^ (see Supplemental Methods for details).

### Illumina microarray data and RNA sequencing (RNAseq)

Illumina microarray data and RNAseq data were performed at the Microarray Core facility at Sanford-Burnham Prebys Medical discovery Institute (SBP) and Genomic Core facility at Indiana University respectively. The standard analyses for microarray^27^ and RNA seq data were performed by the Core facility at each institution (see Supplemental Methods for details).

### Statistical Analysis

Data were presented as the mean ± standard deviation (SD). All experiments were performed with ≥ 3 independent biological replicates (independent experiments) and N represents the number of biological replicates except for cell imaging analysis. For cell imaging analysis, total numbers of counted cells from 3-5 biological replicates were shown unless indicated otherwise. Using the StatView/JMP program (SAS institute, Cary, NC) or GraphPad Prism, statistical difference was analyzed by an ANOVA procedure with post-hoc Tukey/Kramer test for multiple comparisons (labelled as ANOVA), and by the Student’s *t* or Mann-Whitney U test according to the data distribution for pair-wise comparisons. A difference with a p-value < 0.05 was considered statistically significant and was labelled with an asterisk (*) in all figures.

## Results

### Characterization of iPSCs from ACM patients with *DSP* mutations

DSP is a cytoskeletal linker protein with three distinct domains (Figure 1A): 1) the N-terminal plakin domain containing 6 spectrin-like repeats that mediate desmosome formation; 2) the middle coiled-coil rod domain required for DSP homodimerization; and 3) the C-terminal region containing three plakin repeat domains (PRDs) that interact with intermediate filaments such as VIM in MSCs.^29,30^ We first generated patient-specific iPSCs from peripheral blood mononuclear cells (PBMCs) of two unrelated ACM patients with a heterozygous nonsense mutation [*DSP* c.478 C>T (p. Arg160X)] or a heterozygous insertion of an adenine nucleotide that results in frame-shifted C-terminal with premature stop codons [*DSP* c.3474_3475insA (p. Glu1159ArgfsX3)]. We used the genome non-integrating episomal vectors^27^ containing *Oct4*, *Sox2*, *Klf4* and *cMyc* to generate two groups of *DSP*-mutant iPSC lines, and termed here R160X and E1159R lines, respectively for brevity. R160X and E1159R lines are color-coded in blue and green respectively in all figures to facilitate the readability of this paper. Characterization of these iPSCs (Figure S1) shows that all iPSC lines 1) express pluripotent markers including TRA-1-60, TRA-1-81, Nanog, Oct4, and SSEA4 by immunohistochemistry, 2) can differentiate into cells of three germ layers [including ectoderm (Tuj1+), mesoderm (GATA4+), and endoderm (Sox17+)], 3) contain corresponding *DSP* mutations (by genomic DNA sequencing), 4) have normal karyotypes, and 5) exhibit no integration of episomal vectors [verified by qPCR for the exogenous episomal *EBNA-1* and endogenous *FBXO15* expression ratios]. These results demonstrate that we have successfully generated two sets of integration-free iPSC lines with DSP R160X or E1159R truncation mutations (with 3-4 different subclones for each *DSP*-mutant iPSC line, Figure S1F).

**Figure 1.**
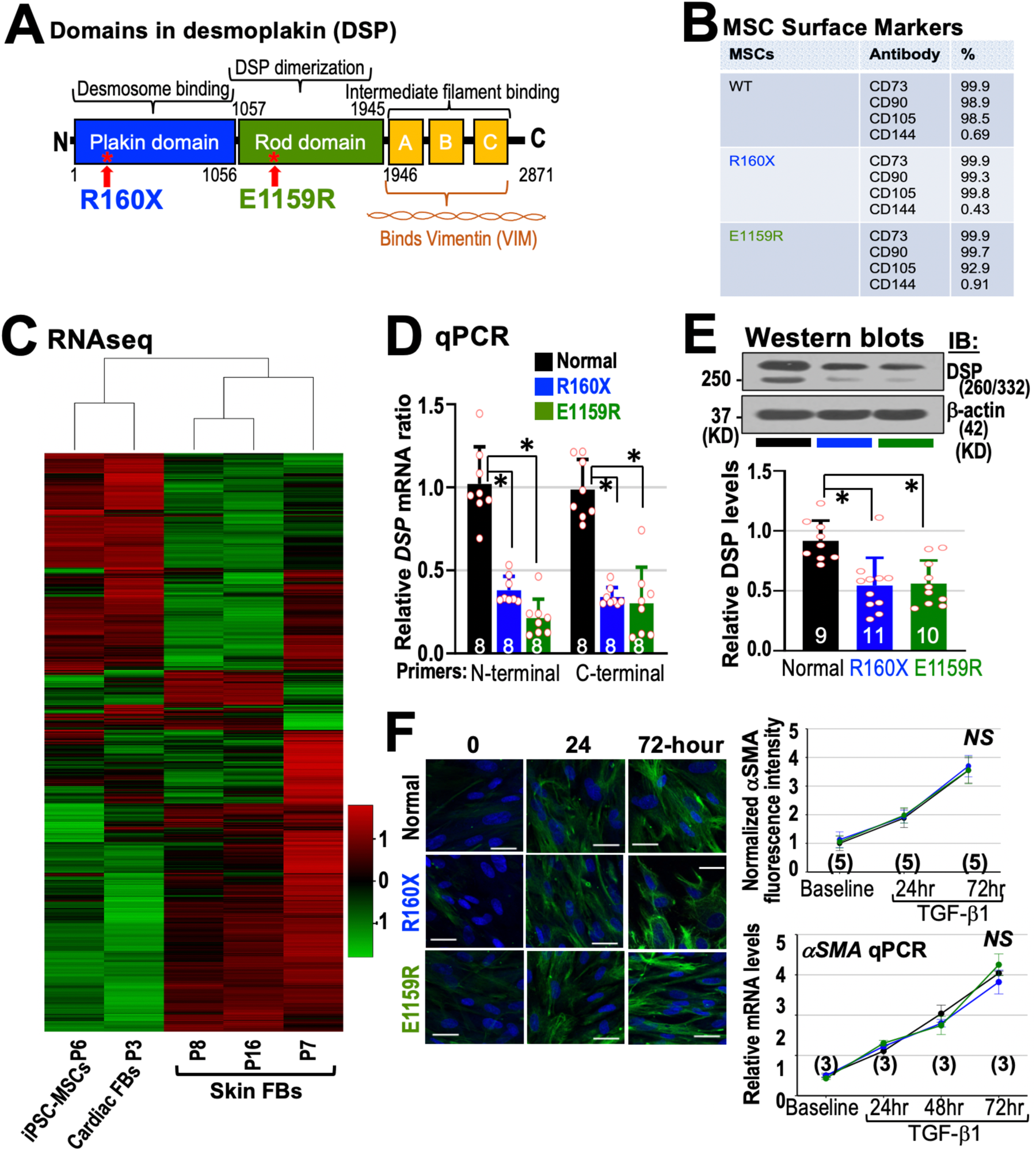
Characterization of iPSC-MSCs. **A**, A simple diagram of 3 structural domains in DSP. **B,** All MSCs express MSC phenotypes by flow cytometry. **C,** An unbiased clustering analysis of global transcriptome mRNA expression profiles (a Heat map) of iPSC-MSCs, cardiac FBs, and skin FBs from the same donor (SW) by RNAseq. Increased and decreased gene expressions are shown in red and green, respectively. **D-E,** *DSP*-mutant MSCs show ≤ 50% WT *DSP* mRNA and DSP protein levels when compared to normal MSCs. **F,** All MSCs expressed similar immunostaining intensities and mRNA levels of aSMA at baseline and after TGFb1. The number of independent biological replicates for each data point is shown in all Figures. Asterisks denote *p* <0.05 by ANOVA except **F** by Kruskal-Wallis. Values are means ± SD. Scale bars are 50 µm. Raw data are in the Source Data Files for verification.

### Characterization of MSCs derived from normal and *DSP*-mutant iPSCs

After successfully generating two different *DSP*-mutant iPSC lines, we used an established protocol to derive MSCs from human iPSCs (iPSC-MSCs) via an early mesoderm intermediate stage as previously described^19^ to simulate the mesodermal origin of cardiac MSCs/fibroblasts.^31^ Because fibroblasts lack internationally-recognized markers for identification^14,15^ and to ensure reproducibility, we used cardiac fibroblast-like iPSC-MSCs as the cardiac fibroblast source for our study. All iPSC-MSCs displayed morphologies indistinguishable from fibroblasts and adhered to standard plastic culture surfaces (Figure S2A). Based on the MSC identification criteria published by the International Society for Cellular Therapy,^16^ flow cytometric analysis revealed high expression of MSC markers (CD73+ = 99.9%, CD90+ ≥ 98.9%, and CD105+ ≥ 92.9%), and low expression of an endothelial marker (CD144+ ≤ 0.91%) in all iPSC-MSCs (Figure 1B and cytometry plots in Figure S2B). A previously published normal iPSC line (SW line)^27^ was used to derive normal iPSC-MSCs. We used the term “Normal” fibroblasts/MSCs throughout this study to describe fibroblasts/MSCs derived or obtained from human donors without any cardiac disease phenotype. To determine if our iPSC-derived MSCs resemble cardiac fibroblasts (FBs) directly sampled from the same iPSC donor in global genomic expression profiles, we obtained FBs directly grown from discarded heart tissues and skin samples of the same normal donor (SW) for comparison. Using RNAseq and unsupervised hierarchical clustering analysis (see Supplemental Methods), the whole genome-wide expression profiles of our iPSC-MSCs are clustered together with cardiac FBs but not skin FBs (at passages 7, 8, and 16; Figure 1C). Thus, instead of using a selected but limited set of markers, we verify the cardiac FB-like nature of our iPSC-MSCs by their close similarity in the whole genome-wide expression profiles to those in cardiac FBs directly sampled from the same donor heart. Thus, we have derived cardiac FB-like MSCs from iPSCs with the advantage that iPSCs could provide limitless supplies of cardiac FB-like MSCs.

Next, we measured WT and mutant *DSP* mRNA and DSP protein levels to functionally characterize these two *DSP*-mutant MSCs lines. We used two qPCR primer sets targeting either N-terminal (113-217bp) or C-terminal (5202-5386bp) regions of WT *DSP* mRNA and found that WT *DSP* mRNA levels of both *DSP*-mutant MSC lines are < 50% of those in normal (SW) iPSC-MSCs (Figure 1D). We used two different DSP antibodies to detect truncated mutant DSP peptides if they were theoretically translated from R160X or E1159R mutations: 1) The DSP antibody 1-100 recognizes the first 1-100 amino acids (a. a.) of DSP and it should detect WT DSP and shorter R160X/E1159R peptides; 2) The DSP 400-500 antibody recognizes the 400^th^-500^th^ a.a. of DSP and it should detect WT DSP and the short E1159R but not R160X peptides (Figure S2C). Western blot (Western) analyses with these 2 DSP antibodies showed that neither *DSP*-mutant MSCs expressed truncated mutant DSP proteins (Figure S2D), but both mutant lines expressed ∼50% of WT DSP protein levels when compared to normal MSCs (Figure 1E). All MSCs expressed similar levels of alpha-smooth muscle actin (aSMA) by immunostaining or by qPCR (Figure 1F) at baseline or after TGFβ1 treatment. These results support that both pathogenic *DSP* variants result in no mutant DSP proteins and a ∼50% reduction of WT DSP protein levels (a common haploinsufficiency phenotype) with similar aSMA expression levels when compared to normal MSCs.

### Increased collagen and vimentin levels in *DSP*-mutant MSCs by TGFβ1

Because TGFβ1 level is elevated in hearts of an ACM mouse model^21^ and in ACM patients,^22^ we first tested fibrogenic responses to 1 ng/ml TGFβ1 in all iPSC-MSCs. In normal MSCs, TGFβ1 should activate fibrogenic gene/protein production within hours and initiate autophagic degradation of collagens later (at ∼24 hours) so as to regulate the fibrogenic response and avoid excessive collagen accumulation^23–25^ (Figure 2A). In culture media, 72 hours (72-hr) of TGFβ1 increased collagen content by 1.52 µg/ml (between black dotted and dashed lines) in normal MSCs with the Sircol Assay, yet 72-hr TGFβ1 increased collagen content by 3.86-4.93 µg/ml in *DSP*-mutant MSCs (between red dotted and black dash lines, representing 2-3-fold increases when compared to the 1.52 µg/ml increase in normal MSCs, Figure 2B). 72-hr TGFβ1 also induced intense intracellular immunostaining of fibrillar collagens (Figure 2C, left image panels) in *DSP*-mutant MSCs but not in normal MSCs. This excessive accumulation of collagens induced by TGFβ1 could be suppressed by various autophagy activators,^32,33^ including 10 nM rapamycin [via mammalian target of rapamycin (mTOR) inhibition], 2.5 µM MK-2206 (via inhibition of Akt1/2/3), and 10 µM trifluoperazine (TFP, an mTOR-independent autophagy inducer) (Figure 2B-C and Figure S3A-B), suggesting that the common autophagic pathway for collagen degradation might be down-regulated in *DSP*-mutant MSCs. The increases of collagen 1 protein (COL1) levels by Western were also about 2-fold higher in mutant than normal MSCs after 72-hr TGFβ1 (Figure 2D). For COL3, the levels were upregulated at baseline and after 72-hr TGFβ1 in both *DSP* mutant iPSC-MSCs when compared to normal MSCs. VIM protein levels [a mesenchymal marker for activated fibroblasts or myofibroblasts (MBs)]^34^ by Western or immune-staining were also increased by > 2 folds in *DS*P-mutant MSCs after 72-hr TGFβ1 when compared to normal MSCs (Figure 2E-F and images in Figure S3C). Collectively, these data indicate that *DS*P-mutant MSCs display > 2-fold increases in fibrogenic responses to TGFβ1 versus normal MSCs.

**Figure 2.**
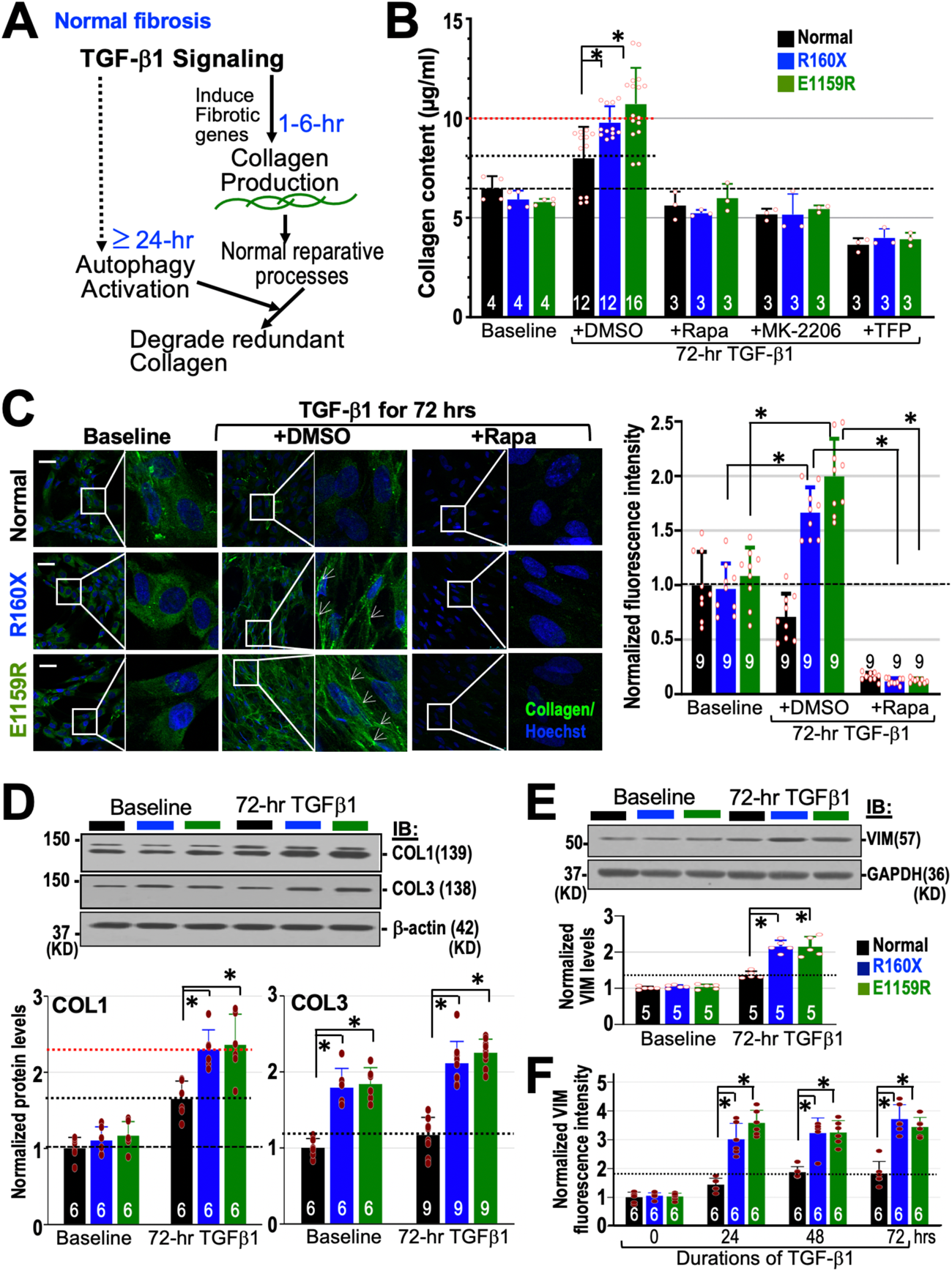
Exaggerated fibrogenic responses to TGFβ1 in *DSP*-mutant iPSC-MSCs. **A**, A controlled reparative TGFβ1 response scheme in normal MSCs. **B,** After 72-hr TGFβ1, the increases in collagen content in culture media were >2-fold more in mutant MSCs than in normal MSCs. **C,** *DSP* mutant MSCs displayed intense fibrillar collagens (white arrows) immunostaining (green) versus normal MSCs after 72-hr TGFβ1, which could be inhibited by 3 different autophagy activators. **D,** Western blots (normalized to normal MSCs at baseline) of fibrotic markers showed higher levels of COL1 (left panel) and COL3 (right panel) in mutant than normal MSCs. **E**, VIM protein levels by Western or (**F**) by immunofluorescent assays also increased by > 2 folds in *DSP* mutant MSCs when compared to normal MSCs after 72-hr TGFβ1. Asterisks denote *p* <0.05 by ANOVA. Scale bars are 50 µm.

### Defective autophagy pathway in *DSP*-mutant MSCs

To further determine if the overall degree of autophagy is decreased in *DSP*-mutant MSCs, we first performed autophagy fluorescent flux assay using the Premo™ Autophagy Tandem Sensor RFP (red fluorescent protein)-GFP (green fluorescent protein)-LC3B Kit (Figure 3A)^35–37^ to screen if the capacity to form autophagosomes and autophagolysosomes in *DSP*-mutant MSCs are decreased before and after TGFβ1 exposure. In this autophagy monitoring system, RFP-GFP-LC3B sensors are activated by the BECN1-containing PI3 kinase complexes and bind to phagophores to form the autophagosomes (positive for both RFP and GFP). Then, autophagosomes would continue to fuse with lysosomes to form autophagolysosome where the acidic environment in lysosomes quenches the green GFP fluorescence and the sensor turns to red color only. Autophagic flux assays indicated that both autophagosome (both RFP+ and GFP+) and autophagolysosome (red dot only) counts were decreased by 40-50% in both *DSP*-mutant MSCs versus normal MSCs at baseline and after 72-hr TGFβ1 (Figure 3B-C), which results are consistent with a decreased overall autophagy flux in *DSP*-mutant MSCs. We then tracked back along the autophagy pathway to find the earliest defective autophagy step in *DSP*-mutant MSCs. We first investigated the LC3 lipidation step that drove phagophores to form autophagosomes. We found that LC3B-I to -II conversion (= LC3B-II/I ratio) was also decreased by ∼50% at baseline and after exposure to TGF-β1 for 24-72 hours in *DSP*-mutant MSCs (Figure 3D), consistent with the results detected by the RFP-GFP-LC3B sensors. During autophagy initiation, BECN1 forms a protein complex with VPS34/15 (subunits of a class III PI3 kinase) and ATG14L (Autophagy Related Gene 14L) (Figure 3A, the color-coded structure at left) that drives LC3B-I to -II conversion via a series of ATG proteins to initiate phagophore elongation and transformation to the autophagosome in mammalian cells.^35–37^ Thus, we examined if BECN1 protein levels were lower in *DSP*-mutant MSCs. Interestingly, we found that total BECN1 protein levels were similar in normal and *DSP*-mutant MSCs at baseline and after exposure to TGFβ1 for 24-72 hours (Figure 3E). Taken together, these results narrowed down the earliest impaired autophagic step(s) in *DSP*-mutant MSCs to processes between BECN1-ATG14L-VPS34/15 PI3 kinase complexes and LC3B-I to -II conversion. Of note, combining all data in Figure 3B-D (and in Figure 4), results from the autophagy fluorescent flux assay are not due to any significant difference in cell sizes before and after TGFβ1 (also see Figure S4A).

**Figure 3.**
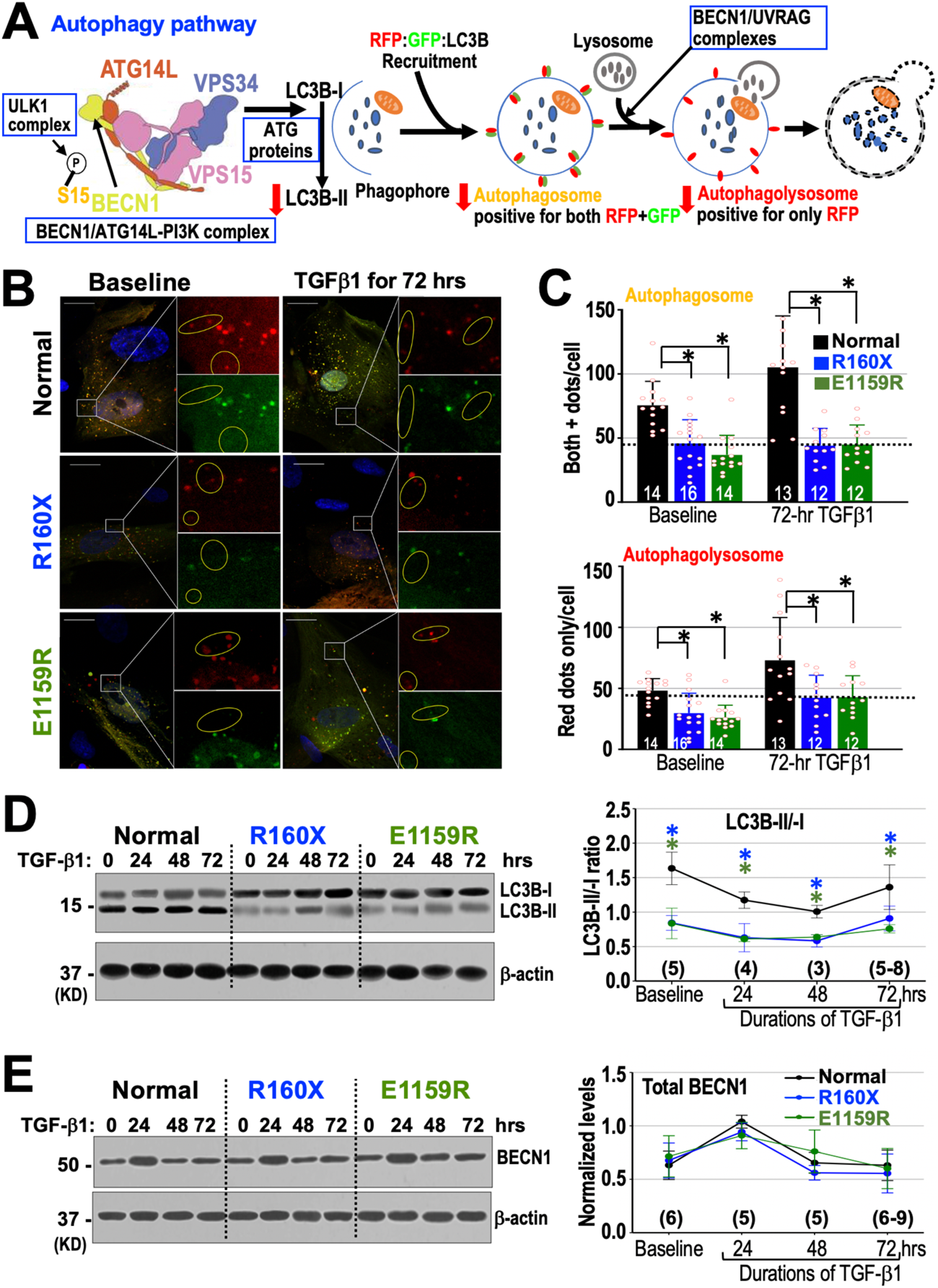
The autophagy deficit in *DSP-*mutant MSCs is located to steps between BECN1 and LC3B-I to –II conversions. **A**, A diagram of the autophagy fluorescent assay using LC3 tandem plasmids to detect autophagosomes (positive for both RFP and GFP) and autophagolysosomes (red dots only, yellow circles in **B**). The BECN1/ATG14L-VPS34/VPS15 complex is color-coded for each component in the model. **B-C,** *DSP*-mutant MSCs showed decreased autophagosomes and auto-phagolysosomes at baseline and after 72-hr TGFβ1 vs. normal MSCs. Scale bars are 25 µm. **D**, LC3B-II to -I ratios at any time points were lower in *DSP*-mutant MSCs, but **E**, total BECN1 protein levels were similar at any time points. * p<0.05 by ANOVA.

**Figure 4.**
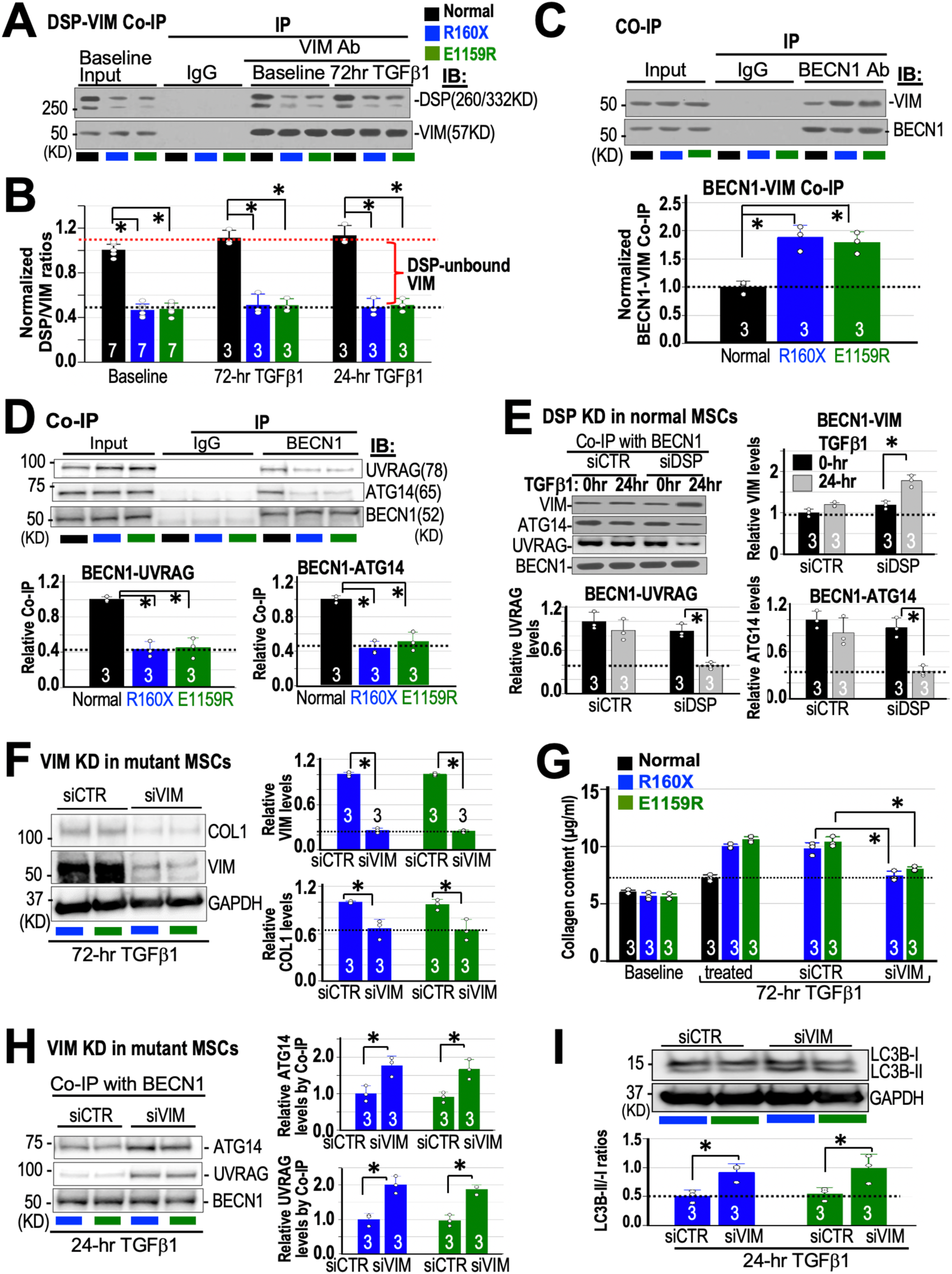
*DSP*-mutant MSCs have less DSP-VIM but enhanced BECN1-VIM associations, leading to decreased BECN1 availability. **A-B**, Using Co-IP, only ∼50% of VIMs were associated with DSP in *DSP*-mutant MSCs when compared to normal MSCs (24-hr TGFβ1 gels are in Figure S4E). **C**, At 24-hrs after TGFβ1, BECN1-VIM associations were ∼2-fold more in mutant MSCs than normal MSCs. **D,** At 24-hrs after TGFβ1, BECN1-ATG14 and BECN1-UVRAG associations in *DSP*-mutant MSCs were ∼50% of those in normal MSCs. **E**, *DSP* KD by siDSP, but not by siCTR in normal MSCs, enhanced VIM-BECN1 binding and decreased BECN1-ATG14 and BECN1-UVRAG associations after 24-hr TGFβ1. **F**, *VIM* KD in mutant MSCs decreased VIM and COL1 levels after 72-hr TGFβ1. **G,** *VIM* KD in both *DSP-*mutant MSCs decreased collagen content in media after 72-hr TGFβ1.

### Increased DSP-unbound VIM restricted BECN1 availability for autophagy in mutant MSCs

Next, we sought to determine whether *DSP* haploinsufficiency led to autophagy defects in stages between BECN1-PI3 kinase complexes and LC3B-I to -II conversion. There are three major known protein complexes between BECN1 and LC3B-I to -II conversion:^35–37^ BECN1-PI3 kinase, a Unc-51 Like Autophagy Activating Kinase 1 (ULK1) complex, and a series of ATG protein complexes (Figure 3A). Since 1) MSCs do not form desmosomes, 2) DSP proteins are not known to interact with ATG proteins, and 3) both DSP^30^ and BECN1^38^ have been shown to bind VIM (the main type of intermediate filaments in MSCs), we hypothesized that DSP might compete with BECN1 for VIM binding. Using Co-IP, we found that VIM-DSP interaction was also reduced by ∼50% at baseline and after 24-or 72-hr TGFβ1 (Figure 4A and 4B) in *DSP-* mutant MSCs. By VIM and DSP immunostaining overlap measurement (Figure S4B-D), the percentage of colocalization of VIM and DSP-stained areas was decreased, resulting in ∼ 2-fold increase of non-DSP overlapped VIMs in *DSP-*mutant MSCs at baseline and after 24-hr TGFβ1 (Figure S4D). We then checked the VIM-BECN1 association after 24-hr TGFβ1 because total BECN1 level reached its peak at 24 hours after TGFβ1 (Figure 3E), which would provide the highest BECN1 protein levels for western blots. With ∼50% less WT DSP and ∼2-fold increased DSP-unbound VIM, the association of BECN1 with VIM by Co-IP increased by ∼2-fold (Figure 4C) after 24-hr TGFβ1 in mutant MSCs. To further support the decreased availability of BECN1 caused by excessive DSP-unbound VIM in *DSP*-mutant MSCs, we checked the levels of association of BECN1 with two known partners in the autophagy and lysosome degradation pathway, ATG14L and UV Radiation Resistance Associated Gene proteins (UVRAG) (Figure 4D)^37^ by Co-IP, respectively. Both ATG14L and UVRAG protein associations with BECN1 were also decreased by ∼50%. These data in Figure 4A-D support strongly that excessive DSP-unbound VIMs actively sequester BECN1 from participating in multiple BECN1-mediated signaling pathways in *DSP*-mutant MSCs (see more in Figures 5-6). To further support the notion that DSP competes with BECN1 for VIM binding, we knocked down *DSP* in normal MSCs with specific siRNA against *DSP* (siDSP) and found that BECN1-VIM association increased, and BECN1 associations with ATG14 and UVRAG decreased accordingly (Figure 4E). The *DSP* knockdown (KD) protocol is shown in Figure S5A. Both *DSP* mRNA and proteins levels were decreased significantly by siDSP when compared to normal MSCs treated with non-silencing control siRNA (siCTR, Figure S5B). To further show that VIM plays an active role in pathological fibrogenesis in DSP-deficient MSCs, we knocked down *VIM* in *DSP-*mutant MSCs with four known siRNAs against *VIM* (siVIM) and found that collagen 1 levels by Western (Figure 4F) and collagen content in media (Figure 4G) decreased to the levels of 72-hr TGFβ1-treated normal MSCs. After *VIM* KD in *DSP*-mutant MSCs, BECN1 associations with ATG14L and UVRAG increased by almost 2-fold (Figure 4H), and LC3B-II to -I ratios also almost doubled (Figure 4I) when compared to MSCs treated with siCTR and TGFβ1 at 24 hours. Taken together, data in Figure 4 support strongly that increased DSP-unbound VIMs sequester BECN1 from actively participating in autophagic degradations of collagens in *DSP* mutant MSCs after TGFβ1. VIM-DSP interactions regulate BECN availability and mediate TGFβ1-induced pathological fibrosis in *DSP*-mutant MSCs. Withaferin A (WFA) is presumed to be an inhibitor of VIM assembly that could decrease fibrotic lung injury at 0.25-1 µM.^39^ Therefore, we explored the potential anti-fibrotic effect of WFA on our MSCs, but WFA, when co-applied with TGFβ1, killed all human MSCs at concentrations as low as 10 nM in one day and could not be tested for its efficacy in reducing pathological fibrosis in *DSP*-mutant MSCs.

**Figure 5.**
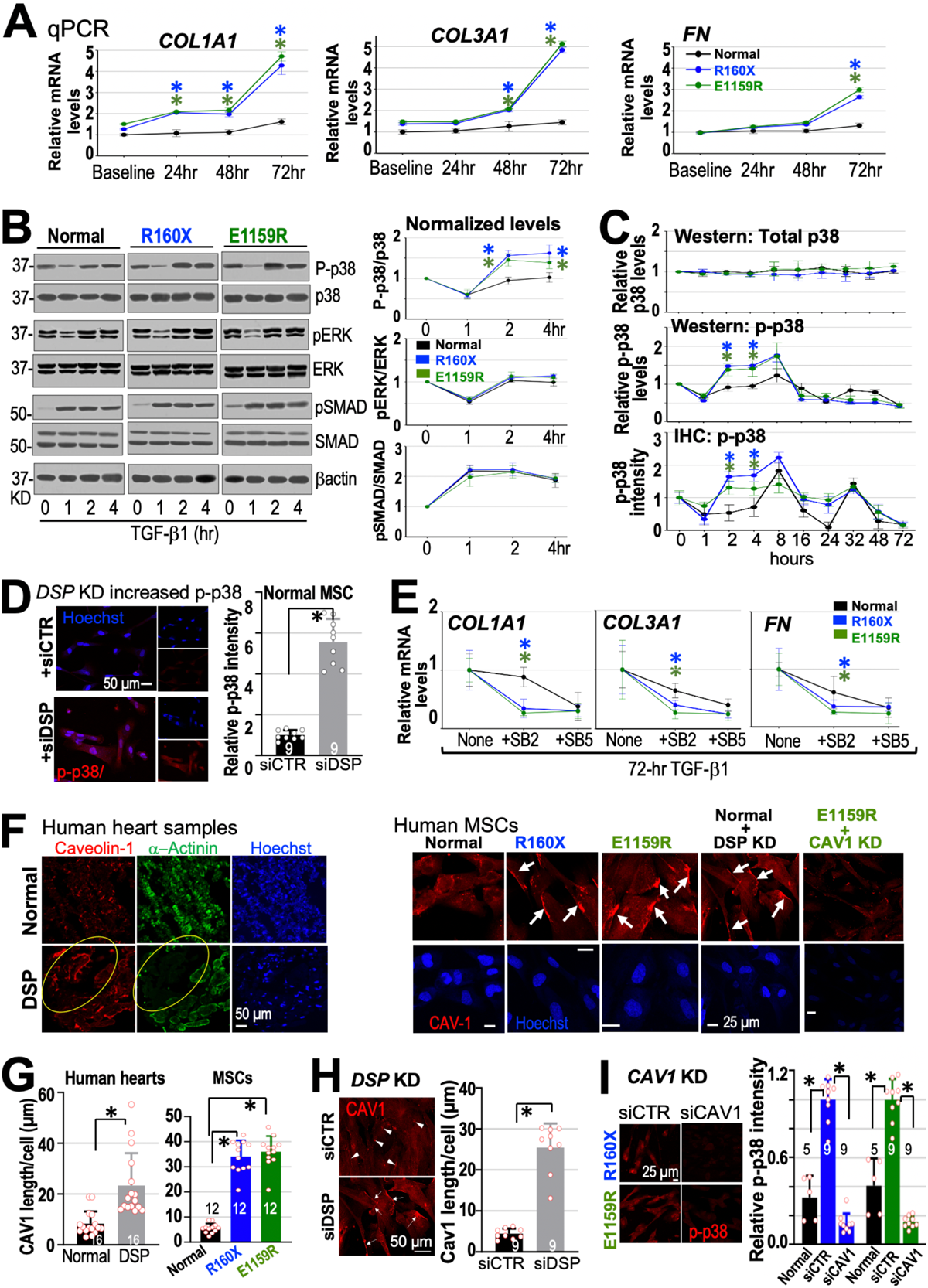
TGFβ1 induced higher fibrotic gene expressions via long CAV1 plaques and p-p38 overactivation in *DSP-*mutant MSCs. **A**, *COL1A1*, *COL3A1*, and *FN* expressions were higher in mutant MSCs (n=3) than normal MSCs (n = 6) after TGFβ1 for indicated durations. **B,** Protein phosphorylation screens showed that only p-p38 pathway was enhanced after TGFβ1 in mutant MSCs (n =5). **C,** Western blots showed (n =3) that total p38 levels (Top panel) did not change with time after TGFβ1 yet p-p38 levels increased in mutant MSCs (mid panel), which could be quickly screened by measuring p-p38 intensity using IHC (bottom panel, n =5). Time scales are for all plots in (**C**). **D,** *DSP* KD in normal MSCs enhanced the p-p38 response to TGFβ1 at 2 hrs. **E,** Only the increased fibrotic gene expressions in mutant MSCs by TGFβ1 could be effectively inhibited by SB2 and SB5 (n =6-12). **F-G,** In human *DSP* mutant heart tissues (yellow circles) and *DSP*-mutant MSCs (arrows), CAV1 formed long plaques when compared to normal counterparts. **H,** *DSP* KD in normal MSCs with siDSP, but not control siCTR (arrowhead), induced long CAV1 plaques (arrows). **I,** *CAV1* KD in *DSP*-mutant MSCs reduced p-p38 responses to 2-hr TGFβ1. * p<0.05 (ANOVA). Scales varied and were shown in images. Number of counted cells from 3 replicates is in **G**-**H**.

**Figure 6.**
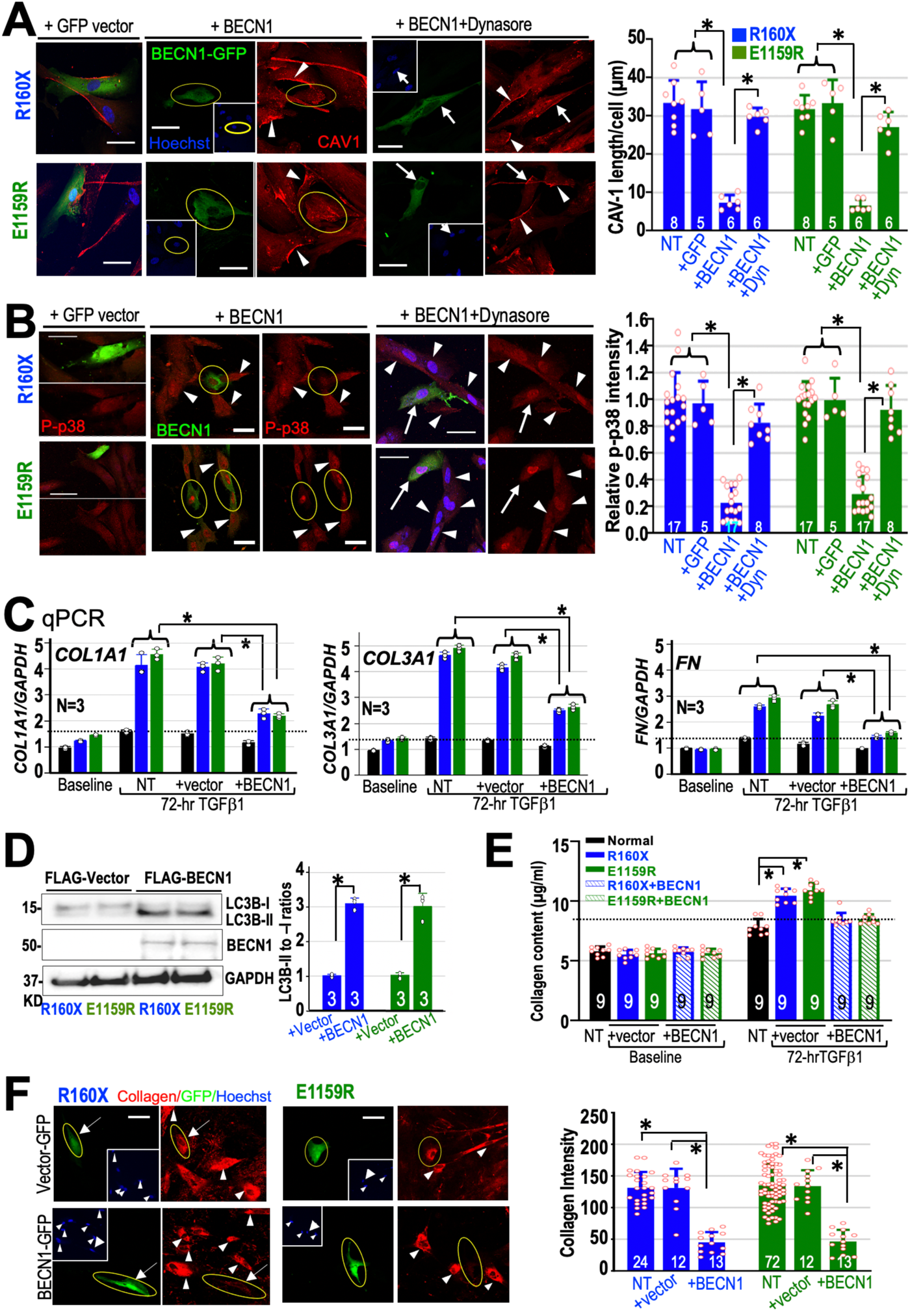
*BECN1* OE reduced long CAV1 plaques, p-p38 overactivation, fibrotic gene expressions, autophagy defects, and collagen accumulations in *DSP-*mutant MSCs. **A**, In *DSP*-mutant MSCs, OE of *BECN1* (yellow circles), but not non-transfected cells (NT, arrowheads) or GFP vectors, reduced long CAV1 plaques at baseline and **B**, p-p38 overactivation by 2-hr TGFβ1, which could be prevented by 1 µM dynasore (Dyn, arrows). **C**, *BECN1* OE, but not NT or vector-transfection (+vector), decreased fibrotic gene expression by TGFβ1. D, *BECN1* OE, but not +vector, enhanced LC3B-I to II conversions. E-F, *BECN1* OE but not +vector or NT (arrowheads), decreased collagens in culture media and collagen staining by 72-hr TGFβ1. * p< 0.05 (ANOVA). The total numbers of counted cells from 3 biological replicates are shown in (**A**), (**B**) and (**F**). For (**C-E**), the numbers of biological replicates are shown. Scale bars are 50 µm.

### Increased fibrotic gene expressions in *DSP-*mutant MSCs via p38 pathway

In addition to activating autophagy, TGFβ1 activates profibrotic/fibrogenic genes during the normal controlled reparative fibrosis.^23,24^ We therefore examined if DSP haploinsufficiency in *DSP*-mutant MSCs also modified expressions of extracellular matrix (ECM) mRNA, such as collagen types 1a1 and 3a1 (*COL1A1 and COL3A1),* and fibronectin (*FN*). We collected mRNAs at baseline, 24, 48, and 72 hours after treating a normal (SW line) and *DSP*-mutant MSCs with TGFβ1, and then performed qPCR to evaluate their fibrogenic gene expressions. We found that levels of *COL1A1, COL3A1,* and *FN* mRNA were significantly higher in both *DSP*-mutant MSCs than those in normal MSCs after 72-hr TGFβ1 (Figure 5A). To further support the hypothesis that DSP plays a key role in increased fibrogenic gene expressions, we showed that *DSP* KD in normal MSCs by siDSP, but not by control nonsilencing siRNA (siCTR), increased expressions of *COL1A1, COL3A1,* and *FN* as well as collagen contents in culture media after 72-hr TGFβ1 (Figure S5C-D). Thus, the increased fibrotic gene expressions and collagen contents by TGFβ1 are specific for DSP-deficient MSCs.

To further understand how DSP deficiency can increase fibrogenic gene expressions, we looked into 3 major TGFβ1-mediated signaling pathways. For fibrogenesis, TGFβ1 activates SMAD2/3 and non-SMAD dependent signaling cascades [p38 and ERK mitogen-activated protein kinase (MAPK) pathways] to induce fibrotic gene expressions in various tissues/organs.^23^ Thus, we explored how DSP haploinsufficiency in MSCs affected the activation of SMAD2/3 and non-SMAD-dependent signaling (p38 and ERK) pathways. Because TGFβ1 usually activates these signaling pathways in timescales of minutes to hours,^40^ we checked the ratios of phosphorylated to total protein levels of SMAD2/3, p38 and ERK1/2 at baseline, 1, 2, and 4 hours after treating MSCs with 1 ng/mL TGFβ1. Western blot analyses showed that only phospho-p38 (p-p38)/p38 ratios in both *DSP*-mutant MSCs were higher than those in normal MSCs at 2-and 4-hr after TGF-β1 (Figure 5B). The phospho-SMAD2/3 (p-SMAD2/3)/SMAD2/3 or phospho-ERK1/2 (p-ERK1/2) to ERK1/2 ratios were similar between normal and *DSP*-mutant MSCs after TGFβ1. We then examined the p-p38 and p38 levels in a longer time frame, since fibrotic genes are significantly elevated at 72 hours after TGFβ1 in *DSP*-mutant MSCs. We found that total p38 levels are relatively unchanged between different MSC lines from baseline to 72 hours after TGFβ1 (Figure 5C, top panel). Changes in p-p38/p38 ratios depended mainly on the p-p38 levels in all MSC lines (Figure 5C, middle panel). Furthermore, in order to quickly screen the potential mechanisms regarding how DSP deficiency enhanced p38 pathway, we measured the intensities of p-p38 immunofluorescence by IHC in MSCs and showed that the changes in p-p38 fluorescent intensities in MSCs reliably mirrored the changes measured by western blots especially in the first 16 hours after TGFβ1 exposure (Figure 5C, lower panel). Moreover, *DSP* KD in normal MSCs enhanced p-p38 levels (Figure 5D), fibrotic gene expressions, and collagen secretions (Figure S5C-D) after TGFβ1, supporting that decreased WT DSP levels are responsible for the enhanced p-p38 and fibrotic responses to TGFβ1. We also used a TGFβ1 receptor blocker [1 µM SB525334 (SB5)] and a p38 inhibitor [1 µM SB202190 (SB2)] to study their effects on the TGFβ1-induced fibrotic gene activation.^41^ As predicted, SB2 inhibited fibrotic gene activations as well as SB5 in *DSP*-mutant MSCs, but TGFβ1 receptor blocker (SB5) is more effective in blocking TGFβ1-induced activations of fibrotic genes in normal MSCs (Figure 5E). These results indicate that DSP deficits lead to specifically enhanced p-p38 activities and subsequently increased fibrotic gene expressions in pathological fibrosis.

### DSP deficiency induced long CAV1 plaques and p38 overactivation

Caveolin-1 (CAV1), an integral membrane protein, is known to form caveolae to terminate non-SMAD signaling pathways after TGFβ1 stimulation in MSCs.^23,24^ Since our data showed that the p38 pathway is abnormally activated in *DSP*-mutant MSCs, we hypothesized that CAV1-mediated termination of TGFβ1-induced p-p38 responses might be down-regulated in *DSP*-mutant MSCs. We first immuno-stained a human ACM heart sample with a heterozygous *DSP* c.4531G>T (p.Gln1511X or Q1511X) mutation, and in both *DSP* mutant MSC lines with a CAV1 antibody. CAV1 is expressed in MSCs but not in cardiomyocytes.^42^ Interestingly, we found that CAV1 formed abnormally long CAV1 clusters on the cell membranes (termed CAV1 plaques here) in the non-myocardial regions (negative for a cardiomyocyte-specific marker, a-actinin immunostainings) of human ACM heart samples with the DSP Q1511X mutation (Figure 5F, left panels) and in two *DSP*-mutant MSC lines (Figure 5F, right panels) at baseline when compared to normal counterparts (Figure 5F-G). Knocking down *DSP* in normal MSCs with siDSP also resulted in abnormal formation of long CAV1 plaques (Figure 5F and 5H). More importantly, knockdown *CAV1* in *DSP*-mutant MSCs eliminated the abnormal activation of p38 pathway after 2-hr TGFβ1, indicating that the abnormally long CAV1 plaques mediate the abnormal p38 overactivation after TGFβ1 in *DSP*-mutant MSCs (Figure 5I). The *CAV1* KD efficiency is shown in Figure S5E.

### Decreased BECN1 availability raises fibrotic genes via long CAV1 plaques and p-p38

Since our data indicate that decreased BECN1 availability is the main reason for decreased autophagy/collagen degradation and rapamycin (an autophagy activator) could suppress collagen accumulation in *DSP*-mutant MSCs (Figures 2-4), we tested whether rapamycin could decrease CAV1 plaques. Treatment of *DSP*-mutant MSCs with 2 days of 10 nM rapamycin minimally changed the length of CAV1 plaques in *DSP*-mutant MSCs (Figure S5F). However, BECN1 has been shown to affect various types of endocytosis and endocytic trafficking.^43^ We therefore hypothesized that BECN1 availability might also affect the endocytosis of CAV1 and tested if overexpression (OE) of *BECN1* could mitigate the abnormal CAV1 plaque formation on cell membranes and p-p38 over-activation in *DSP*-mutant MSCs. We found that OE of *BECN1* for 2 days eliminated the abnormally long CAV1 plaques (Figure 6A) and significantly decreased fluorescent intensities of p-p38 at 2 hours after TGFβ1 (Figure 6B) in *DSP*-mutant MSCs. These effects of *BECN1* OE on CAV1 plaques and p-p38 intensity could be blocked by co-treatment with an endocytosis/dynamin inhibitor, Dynasore (Dyn).^44^ OE of *BECN1* (with 50-70 % transfection efficiency by electroporation) also decreased fibrotic gene expression levels (*COL1A1, COL3A1,* and *FN*) at 72 hours after TGFβ1 in *DSP*-mutant MSCs when compared to non-transfected (NT) or vector-transfected MSCs (+vector)(Figure 6C). Taken together, these data support that DSP deficiency in *DSP*-mutant MSCs reduces BECN1 availability for CAV1-mediated endocytosis, leading to long CAV1 plaque formations and subsequent p-p38/fibrotic gene over-activation after TGFβ1. OE of *BECN1* restored LC3B-II/-I ratios in autophagy (Figure 6D), reduced fibrillar collagen accumulation/secretion (Figure 6E), and decreased collagen immune-staining in *DSP-*mutant MSCs after 72-hr TGFβ1 (Figure 6F).

### Over-expressing VIM-binding domains of DSP reduces pathological fibrosis

Exogenous expression of PKP2 proteins has been proposed to treat ACM patients with *PKP2* mutation.^45^ Since DSP proteins exist and interact with vimentin in all fibroblasts, over-expression of *DSP* might suppress all pathological fibrosis by increasing BECN1 availability for greater collagen autophagic degradation and CAV1-p38 pathway termination. However, desmoplakins are large-size proteins and have two isoforms created by alternative splicing: DSP-I (∼332 kD) and DSP-II (∼260 kD).^29,30^ Expressing the full-length *DSP* might not be efficient *in vivo* and would be hard to be used for clinical therapy. We thus subcloned a hemagglutinin (HA)-tagged DSP fragment with deleted N-terminal domain (named HA-DSPΔN containing the minimal required domains for VIM binding)^46^ to perform rescue experiments in MSCs. The HA-tag would allow us to identify exogenous expressed DSPΔN proteins by HA antibody but not by our DSP antibodies against the DSP N-terminal domains. Over-expressing HA-DSPΔN in *DSP* mutant MSCs restored their VIM-DSP associations after TGFβ1 (Figure 7A), LC3B-II/-I ratios at baseline (Figure 7B), collagen contents in culture media (Figure 7C) and collagen IHC staining (Figure 7D) after TGFβ1 to the levels of normal MSCs. Over-expressing HA-DSPΔN in mutant MSCs also decreased BECN1-VIM associations and increased associations of BECN1 with UVRAG and ATG14 (complexes mediating autophagy, Figure 7E). CAV1 plaques lengths, phospho-p38 IHC staining levels, and subsequent fibrotic gene activations (*COL1A1, COL3A1,* and *FN*) after TGFβ1 are all decreased by over-expressing HA-DSPΔN in mutant MSCs (Figure 7F-H). Overall, our results support strongly that *DSP* mutations lead to DSP deficiency, which mediates the pathological fibrogenesis of mutant cardiac MSCs by decreasing BECN1 availability via increased BECN1-VIM interactions. Over-expressing DSPΔN might be a new strategy to combat “pathological” fibrosis in other diseased organs by increasing autophagic degradation of collagens and decreasing CAV1-p38-mediated fibrotic gene activation.

**Figure 7.**
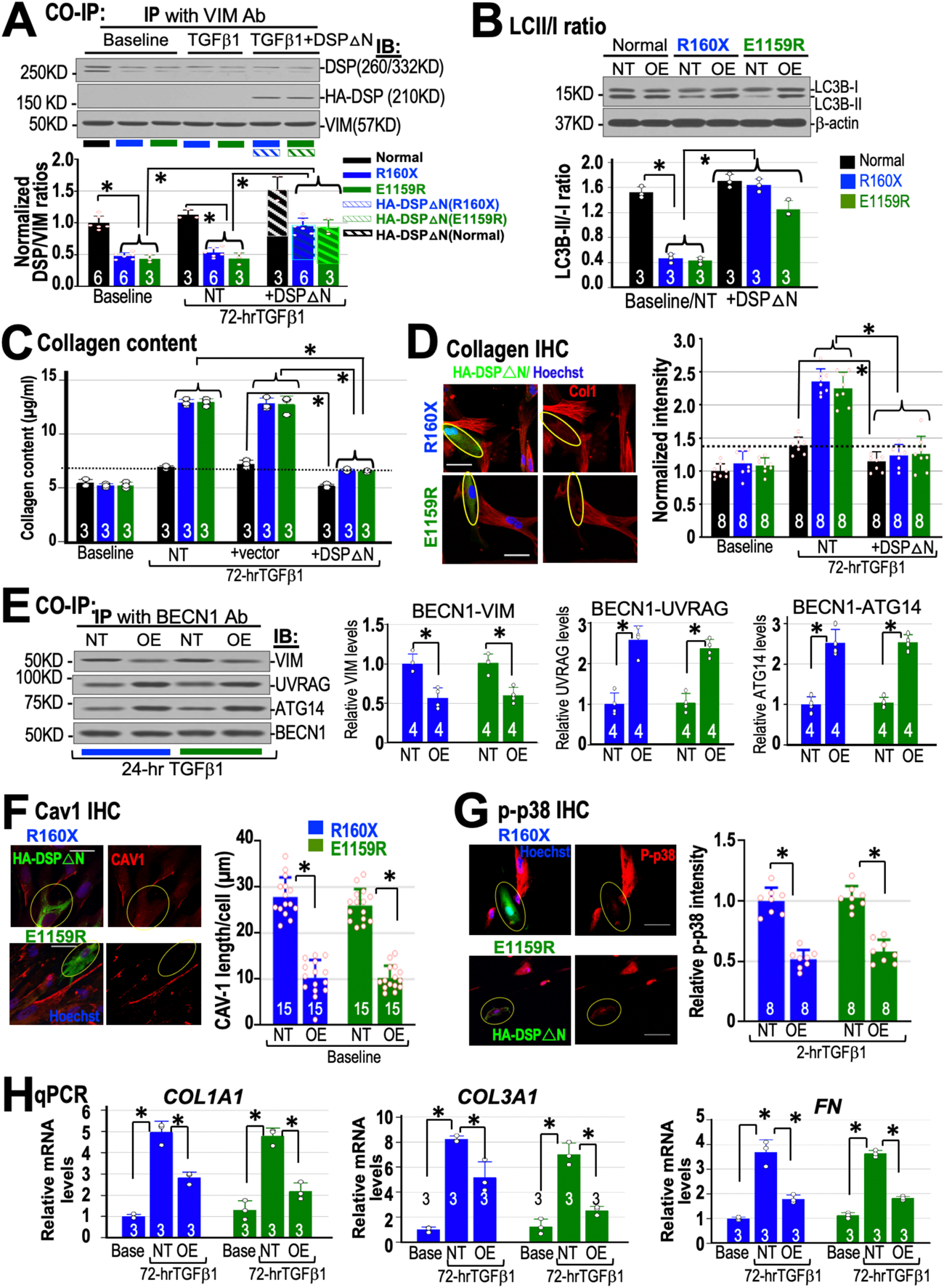
OE of DSPΔN ameliorates the pathological fibrogenesis in mutant MSCs. **A**, DSPΔN OE increased VIM-DSPΔN association (also see Figure S6A) and (**B)** LC3B-II to -I ratios in mutant MSCs. **C**, DSPΔN OE decreased collagen contents and (**D)** collagen staining in mutant and normal MSCs when compared to no (NT) or vector transfection (+vector). **E,** DSPΔN OE decreased VIM-BECN1 but increased BECN1-UVRAG and BECN1-ATG14 associations in mutant MSCs. **F,** DSPΔN OE (yellow circles) decreased CAV1 plaque length at baseline (Base) and (**G)** p-p38 intensities by TGFβ1 in mutant MSCs. **H,** DSPΔN OE decreased fibrotic gene levels by TGFΔ1 when compared to NT cells. * *p* <0.05 by ANOVA (except **B, C, H** by Mann-Whitney). Scale bars= 50 µm.

### Heterozygous DSP deletion in mouse myofibroblasts in vivo

To obtain *in vivo* evidence that DSP deficiency in activated fibroblasts (myofibroblasts) enhances fibrogenic responses to TGFβ1 *in vivo*, we used tamoxifen-controlled, periostin-specific cre recombinase (*POSTN^MRM^*-Cre+)^31^ to heterozygously delete *DSP* in activated fibroblasts of *DSP*-floxed mice (*DSP^w/f^*) with 5-day tamoxifen i. p. injections. 6-week-old mice were i. p. injected with 1 µg TGFβ1 daily for 7-8 days and mouse hearts were then harvested for comparisons at the 7^th^ week. When compared to *POSTN^MCM^*-Cre-negative; *DSP^w/f^* mice, *POSTN^MCM^*-Cre-positive; *DSP^w/f^* mice showed increased cardiac fibrosis (stained by Sirius red, Figure 8A), decreased autophagy (lower LC3B-II/I ratios and higher p62 levels), higher p-p38 and fibrotic gene levels, and lower DSP mRNA/protein levels (Figure 8B-D). Of note, sequestosome 1 (SQSTM1 or p62) binds LC3B to delivery ubiquitinated cargo proteins for autophagic degradation. As such, p62 accumulations with low LC3B-II/I ratios are indicators for suppressed autophagy.^38^ Additionally, when compared to 3 normal human donor hearts, the human ACM heart with a heterozygous Q1511X mutation (from Figure 5F) also showed increased fibrotic gene levels, higher p-p38/p38 ratios, and decreased autophagy markers (lower LC3B-II/I ratios with elevated p62 levels)(Figure S6B-C). Importantly, cardiac fibroblasts directly isolated from *POSTN^MCM^*-Cre-positive mouse hearts showed higher BECN1-VIM associations and lower BECN1-ATG14/BECN1-UVRAG associations (Figure 8E) after TGFβ1. The Cre-positive mouse cardiac fibroblasts also showed long CAV1 plaques at baseline (Figure 8F) and higher collagen secretions and p-p38 levels after TGFβ1 (Figure 8G-H). Taken together, these data strongly support that DSP deficiency led to increased cardiac fibrosis after TGFβ1 *in vivo* via the same pathogenic mechanisms as we showed in Figures 1-7. Most importantly, we co-immunostained human MSCs with CAV1 and type 1 TGFβ receptor (TGFBR1) antibodies. We found that CAV1 plaque-TGFBR1 colocalizations on cell membranes in *DSP* mutant MSCs were higher at baseline, which increased significantly after 2-hr TGFβ1 when compared to normal MSCs (Figure 8I and S6D), supporting strongly that CAV1-mediated TGFBR1 endocytosis^23,24^ was downregulated in mutant MSCs. Figure 8J showed a diagram summarizing the pathogenic network specific to deficient VIM-DSP interactions, which led to decreased BECN1 availability and enhanced fibrogenic responses to TGFβ1 in DSP deficient MSCs.

**Figure 8.**
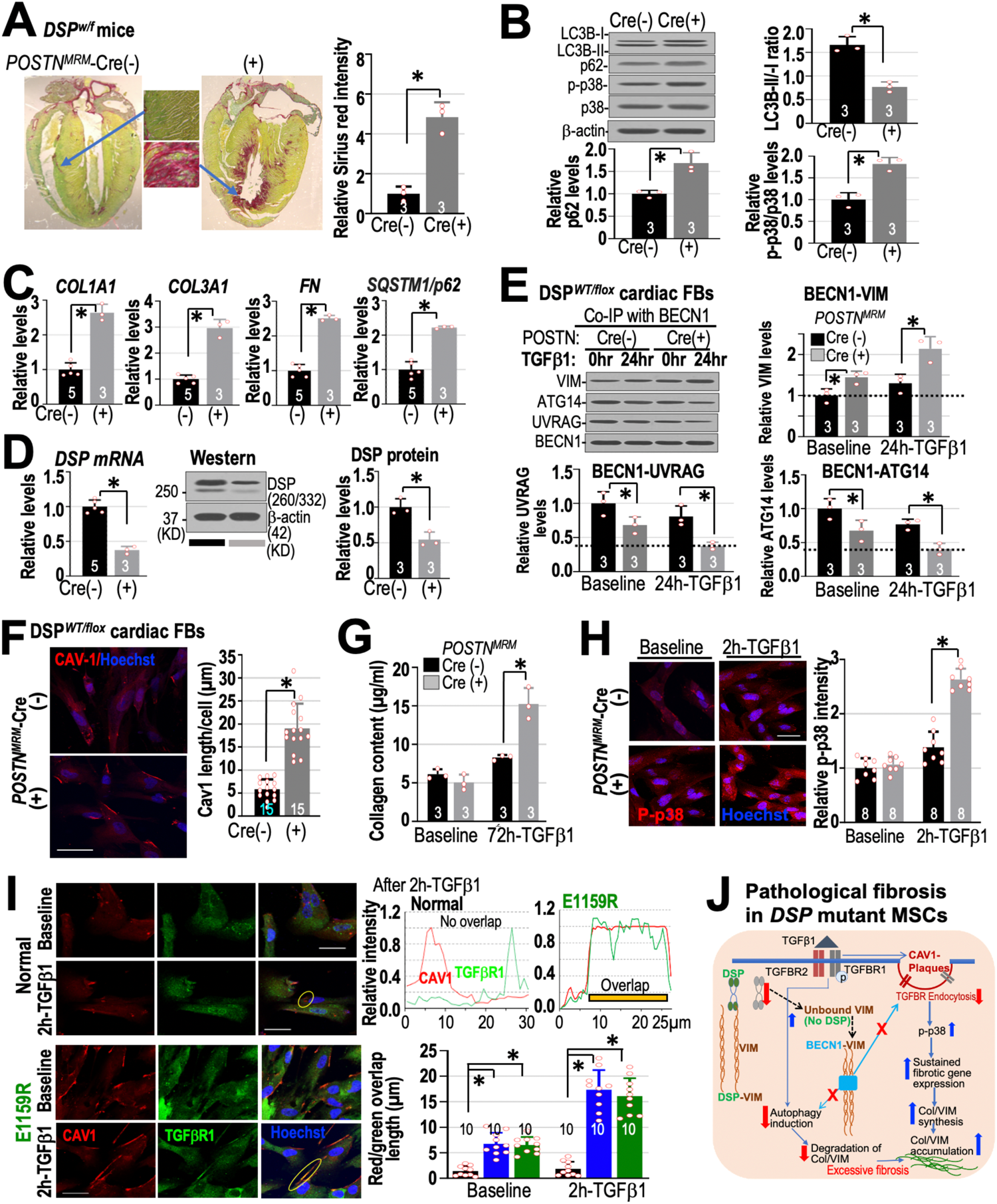
Hearts and cardiac fibroblasts from *DSP^w/f^* mice with *POSTN*^MRM^-Cre-positive or -negative genotypes. *DSP^w/f^* mouse hearts with *POSTN^MRM^*-Cre (Cre+) showed **A**, enhanced fibrosis to TGFβ1 *in vivo* by Sirius red-Fast green staining; **B**, decreased autophagy (low LC3B-II/I ratios and high p62 levels), overactivated p-p38, **C**, higher fibrotic gene expressions and **D,** lower DSP levels when compare to Cre-negative mice. *DSP^w/f^*mouse cardiac fibroblasts with Cre+ showed **E,** higher BECN1-VIM binding but lower BECN1-UVRAG/ BECN1-ATG14 associations; **F,** long CAV1 plaques at baseline, **G,** higher collagen secretion by TGFβ1, and **H,** higher p-p38 levels by TGFβ1. **I,** Using co-immunostaining, CAV1-TGFBR1 colocalizations at cell membrane (yellow circles) were higher at baseline and increased significantly after 2-hr TGFβ1 in human *DSP* mutant MSCs (Figure S6D) but not in normal MSCs. **J**, A simple diagram for TGFβ1-mediated pathological fibrosis due to reduced BECN1 availability in *DSP* mutant MSCs. Scale bars are 50 µm. All data were from 3 independent experiments. * p< 0.05 by Mann-Whitney (except **F, H** and **I** by ANOVA).

### DSP mutations and fibrogenic pathways by gene expression analysis

To obtain a global view of how DSP deficiency affects the gene expression networks (Figures S6E-F and S7), we used Illumina mRNA expression microarrays to profile iPSC-MSCs from DSP R160X proband, DSP R160X proband’s mother (carrying the same heterozygous *DSP* mutation but with no clinical ACM), DSP E1159R proband, and the normal (SW) line at baseline to find common deregulated pathways. Notably, iPSC-MSCs derived from asymptomatic R160X proband’s mother (a mutation carrier) had ∼40% of WT *DSP* mRNA levels but ∼ 82% of WT DSP protein levels (vs. normal MSCs), did not have long CAV1 plaques, did not show increased productions of collagens (1 and 3) and VIM, nor increased fibrotic gene expressions after 72-hr TGFβ1 (Figure S7A-E). To find the common differentially expressed genes between 2 ACM patient MSC lines and 2 MSC lines without clinical ACM, we first generated 3 sets of pairwise comparisons from the mRNA expression profiles: 1) DSP R160X proband vs. his mother (Mother) with no clinical ACM, 2) DSP R160X proband vs. a normal donor without ACM (SW), and 3) DSP E1159R proband vs. the normal SW donor. We then identified the consistently up- or down-regulated (by 1.3-fold) genes common to these 3 pairwise comparisons (total 1186 genes, Figure S6E-F). Using the DAVID 6.7 KEGG pathway analyses, we identified the top 15 dysregulated pathways for the two *DSP*-mutant MSCs with clinical ACM, which showed altered expressions of genes in extracellular matrix (ECM), focal adhesion, several cancer pathways, axon guidance (axon guide), vascular smooth muscle contraction (VSMC), hematopoietic cell lineage, cytokine-cytokine receptor interaction (cytokine interact), viral myocarditis, and ACM (termed ARVC in the KEGG pathway), as well as signaling pathways of TGFβ, Gonadotropin-releasing hormone (GnRH), and Jak-STAT pathways. These results further support our findings that DSP deficiency plays a key role in pathological fibrogenic responses to TGFβ1 in *DSP*-mutant MSCs because ECM, focal adhesion pathway, TGFβ pathway, and ACM are known pathways that are related to pathological cardiac fibrosis in cardiomyopathic hearts.

Of note, we also studied TGFβ1 response of a plakophilin 2 (*PKP2)* haplo-insufficient ACM MSC line without DSP deficiency (Figure S7A-B, the delC line with a *PKP2* c.2013delC mutation that was previously characterized in Ref. 27). When compared to *DSP* mutant MSCs, delC *PKP2*-deficient MSCs did not have excessive productions of collagens 1/ 3 or VIM after 72-hour TGFβ1 (Figure S7B-C), have no abnormally long CAV1 plaques (Figure S7D) nor excessive fibrotic gene expressions (Figure S7E). Thus, the pathogenic mechanisms presented here for increased fibrogenic responses to TGFβ1 are specific for DSP-deficient MSCs but not for PKP2-deficient MSCs.

## Discussion

Pathogenic variants or single nucleotide polymorphisms in the *DSP* gene had been implicated in the pathogenesis of arrhythmogenic cardiomyopathy,^5,6^ IPF,^7^ hypertrophic scars,^8^ and cancer metastasis^47^ with unknown mechanisms. Here, we used patient-specific iPSC-MSCs derived from ACM patients with *DSP* mutations to elucidate the novel pathogenic roles of DSP deficiency in pathological fibrosis. We first showed that 2 different *DSP* truncation mutations in MSCs resulted in no new mutant DSP protein but ∼50% reduction of WT DSP (an DSP haplo-insufficient phenotype, Figure 1). DSP-deficient MSCs displayed exaggerated collagen production/secretion and overactivated fibrotic gene expressions (Figures 2 and 5). Importantly, DSP deficiency resulted in decreased DSP-VIM associations and increased VIM-BECN1 interactions, which restricted BECN1 availability and led to decreased autophagic degradation of collagens due to decreased BECN1-ATG14 and BECN1-UVRAG complexes (Figures 3-4).

Decreased BECN1 availability also induced abnormal formation of long CAV1 plaques that over-activated the p38 pathway after TGFβ1 and increased fibrotic genes and protein levels. These fibrogenic pathologies can be reproduced by *DSP* knockdown in normal MSCs *in vitro* (Figures 5-6 and Figure S5) and in mouse activated fibroblasts *in vivo* (Figure 8). As predicted, over-expression of *BECN1* (Figure 6) or *HA-DSPΔN* (Figure 7) rescued all pathological fibrotic responses to TGFβ1 in *DSP* deficient MSCs. Genome-wide comparisons by unbiased mRNA expression array analysis further supported the role of DSP in pathological cardiac fibrosis in *DSP* deficient MSCs (Figure S6E-F).

Vimentin is a type III cytoskeletal intermediate filament that has been shown to maintain cell structural integrity, organize cytosolic organelles, and participate in cell adhesion, migration, invasion, and signaling.^48^ Structurally, VIM is composed of a head domain that can bind BECN1,^39^ a central coil-coil domain responsible for VIM dimer/tetramer formation that can interact with the C-terminal PRD domains of DSP,^49^ and a C-terminal tail domains. Vimentin is the only cytoplasmic intermediate-filament system in fibroblasts^50^ and has been associated with inflammatory and autoimmune diseases, cancers, and various fibrotic diseases^50^ via unknown mechanisms. Of note, VIM is frequently used as a classic marker for cells of mesenchymal origin, cells undergoing epithelial-to-mesenchymal transition, and activated fibroblasts (myofibroblasts).^34^ Whether VIM actively participates in fibrogenesis remain to be determined. Although VIM has been shown to bind BECN1 and decrease autophagy,^38^ the mechanisms that regulate VIM-BECN1 interactions for fibrogenesis remain unclear. Our study indicates that DSP is the main regulator of BECN1-VIM interaction during fibrogenesis. In DSP deficient MSCs, diminished amounts of WT DSP proteins increase the levels of VIM-BECN1 associations.

Increased VIM-BECN1 binding in turn restricts BECN1 availability for initiating autophagy (by decreased BECN1/ATG14L-PI3K complexes) and for autophagosome-lysosome fusion (by decreased BECN1/UVRAG complexes), leading to decreased collagen degradation and increased cytosolic fibrillar collagen and collagen secretion.

Interestingly, vimentin has been shown to stabilize *COL1A1* mRNA between 6 and 24 hours after TGFβ without affecting fibronectin levels.^51^ This particular mechanism could only explain the *COL1A1* mRNA expression pattern at 24 hours but not at 72 hours after TGFβ1 in our experiments (Figure 5A) because *COL1A1, COL3A1* and *FN* mRNA levels were all significantly upregulated at 72 hours after TGFβ1, which could be effectively inhibited by a p38 blocker (Figure 5E).

Furthermore, knockdown or knockout of *DSP* in mouse cardiomyocytes and epidermal cells was associated with the abnormal up-regulation of TGFβ1/p38 signaling via unknown mechanisms.^52^ Here, we show a novel finding that *DSP* knockdown or DSP deficiency in MSCs induces abnormal formation of CAV1 plaques due to defective endocytosis of CAV1.

Importantly, abnormal CAV1 plaques are the main mechanism for over-activating p-p38 and subsequent fibrotic gene expression (*COL1A1, COL3A1* and *FN*) by TGFβ1. Of note, MSCs derived from an ACM patient with *PKP2* haploinsufficiency (the delC line) did not have DSP deficiency, did not show any abnormal CAV1 plaque, nor enhanced fibrogenic responses to TGFβ1 (Figure S7). These novel findings uncover the missing link between DSP deficiency and exaggerated p38 pathway activation in cardiac fibroblasts/MSCs.

Our data also have significant therapeutic implications for treating pathological fibrosis. Silencing VIM in fibroblasts may lead to defects in wound healing and cellular polarity.^39^ Non-specific autophagy activators could degrade other cellular proteins, leading to unwanted side effects. Downregulating CAV1 remains an uncertain anti-fibrotic strategy because CAV1 has dynamic roles in normal fibrogenic processes.^53^ Drugs that inhibit VIM assembly, such as WFA, had been shown to reduce invasiveness and fibrogenic potentials of fibroblasts in IPF at 0.25 to 1 µM.^39^ However, WFA has pleiotropic effects on various types of cells, including reduction of inflammatory cytokines and attenuating profibrotic gene expressions (including *COL1A2, COL3A1, & FN*),^54^ but can cause cancer cell apoptosis.^55^ These widespread effects of WFA might indicate that WFA could affect fibrosis via mechanisms in addition to just increasing VIM disassembly. In our *DSP* mutant MSC cultures, co-application of WFA even at 10 nM with TGFβ1 killed most cells in one day and thus we could not further study the anti-fibrotic effect of WFA. Importantly, efficient overexpression of a VIM-binding domain of DSP protein may specifically tackle any pathological fibrosis mediated by over-activated VIM-BECN1 interactions. Thus, over-expressing VIM-binding domains of DSP potentially might be used to treat DSP cardiomyopathy and other types of pathological fibrosis with exaggerated VIM expressions. Also, a p38 inhibitor, ARRY-371797, has been used to treat cardiomyopathy from Lamin A/C mutations (NCT03439514 in ClinicalTrials.gov)^56^ and potentially could be used to treat cardiomyopathy with *DSP* mutations or other organ fibrosis due to DSP deficiency. Finally, reduced DSP levels are associated with carcinogenesis.^57^ Our novel findings may instigate research for using VIM-binding domains of DSP as a novel therapy against cancer invasion or metastasis.

## Nonstandard abbreviations and Acronyms

ACM: Arrhythmogenic Cardiomyopathy
iPSC-CM: Induced pluripotent stem cell-derived Cardiomyocyte
MSC: Mesenchymal stromal cell

## Acknowledgments

We thank all patients for their participation, Microarray and Genomic Core facilities at SBP Medical Discovery Institute and at Indiana University for their support, Johnson Wong, Ed Simpsons and Yunlong Liu for RNA seq analysis, Tiara Tirasawasdichai for conducting some preliminary experiments during her graduate studies, and AnaBios Inc. for normal donor heart tissues.

## Author contributions

C.W., S-F.C. and H-S.V. C. designed experiments and wrote the manuscript; C.W., S-F.C. and H-S.V.C. performed experiments and data analysis. A.M.S., C.B., F.D., J.E.M., C.A.J., H.C., W.S., and D.P.J. provided clinical assessment and patient’s fibroblasts/heart tissues as well as helped in preparation of this manuscript. All authors read and approved the manuscript; and H.-S.V.C. supervised the entire research project.

## Supplemental Materials

Supplementary Methods

Supplemental Figures S1–S7

Supplemental Tables S1–S5,

Source data file for Figures 1–8 and S1–S7

Source data file for Figures 1C and S6E-F

Source Figures with unprocessed blots.

## Novelty and Significance

### What Is Known?

- Desmoplakin variants and/or deficiency are linked to several pathological fibrogenic diseases, including desmoplakin cardiomyopathies.
- Cardiac fibroblasts are known sources for cardiac fibrosis but cardiac fibroblasts are a heterogenous group of cells and lack internationally recognized markers for their identifications.

### What New Information Does This Article Contribute?

- We used unbiased genome-wide analyses, rather than selective markers, to generate cardiac fibroblasts-like, induced pluripotent stem cell-derived mesenchymal stromal cells (MSCs) to study pathological fibrosis in desmoplakin cardiomyopathy.
- We show for the first time that both desmoplakin and vimentin play active roles in mediating pathological fibrosis by regulation beclin-1 availability.
- Desmoplakin deficiency decreases collagen autophagic degradation and causes abnormal formation of long CAV1 plaques that in turn over-activates p38 pathway and pro-fibrogenic genes after TGFβ1 *in vitro* and *in vivo*.
- A truncated desmoplakin could increase collagen degradation and decrease profibrotic gene activation, which might be used to treat other pathological fibrosis.

Desmosome gene mutations are linked to several fibrotic diseases, including arrhythmogenic cardiomyopathy (ACM). A better understanding of how *DSP* mutations affect pathological fibrosis might lead to novel therapies. Cardiac fibroblasts were usually identified via selective markers without international consensus. We first use genome-wide expression analyses to generate human cardiac fibroblast-like iPSC-MSCs, which can provide unlimited supplies for pathogenic and therapeutic studies. We then show that *DSP-*mutant cardiac MSCs display exaggerated fibrogenic responses to TGFβ1 with excessive accumulations of vimentin (VIM)/fibrillar collagens and over-activated fibrotic genes. We further find that DSP competes with beclin-1 (BECN1) for their associations with VIMs. Pathogenic DSP variants lead to DSP deficiency and increased VIM-BECN1 binding, which sequester BECN1 from activating autophagy and from inducing caveolin-1 (CAV1)-mediated endocytosis. Decreased autophagy causes collagen accumulations and diminished CAV1 endocytosis results in abnormal CAV1 plaque formation that over-activates fibrogenic genes via heightened p38 activities after TGFβ1. A truncated VIM-binding DSP protein could free up BECN1 to increase collagen autophagic degradation and downregulate fibrotic gene expressions. This truncated DSP transcript might offer a new therapeutic strategy to treat other pathological organ fibrosis.

